# Dynamic and Selective Low-Complexity Domain Interactions Revealed by Live-Cell Single-Molecule Imaging

**DOI:** 10.1101/208710

**Authors:** Shasha Chong, Claire Dugast-Darzacq, Zhe Liu, Peng Dong, Gina M. Dailey, Claudia Cattoglio, Sambashiva Banala, Luke Lavis, Xavier Darzacq, Robert Tjian

## Abstract

Many eukaryotic transcription factors (TFs) contain intrinsically disordered low-complexity domains (LCDs) but how they perform transactivation functions remains unclear. Recent studies report that TF-LCDs can undergo hydrogel formation or liquid-liquid phase separation *in vitro*. Here, live-cell single-molecule imaging reveals that TF-LCDs form local high concentration interaction hubs at synthetic and endogenous genomic loci. TF-LCD hubs stabilize DNA binding, recruit RNA polymerase II (Pol II) and activate transcription. LCD-LCD interactions within hubs are highly dynamic, display selectivity with binding partners, and are differentially sensitive to disruption by hexanediols. These findings suggest that under physiological conditions, rapid reversible and multivalent LCD-LCD interactions occur between TFs and the Pol II machinery, which underpins a central mechanism for transactivation and plays a key role in gene expression and disease.

## Introduction

Sequence specific DNA-binding transcription factors (TFs) are seminal players in eukaryotic gene regulation. From the earliest studies of human TFs it was recognized that regulatory proteins like Sp1 contain well-structured DNA binding domains (DBDs) and functionally critical transactivation domains that participate in specific TF-TF interactions to direct gene transcription *(1–3)*. Numerous atomic structures of DBDs have provided a concrete understanding of TF-DNA interactions. In contrast, many transactivation domains contain low-complexity sequence domains (LCDs) that persist in an intrinsically disordered conformation not amenable to conventional structural determination. Mutations in TF-LCDs not only disrupt transcription but also have been implicated in cancer and neurodegenerative disorders *(4, 5)*. However, how TF LCDs execute specific transactivation functions has remained a long-standing enigma. Equally challenging has been cracking the code for how LCDs operate *in vivo* given the dynamic nature of TF-TF interactions required for gene regulation.

A wealth of *in vitro* studies suggests that purified LCDs from the FET protein family (FUS/EWS/TAF15) can undergo reversible hydrogel formation or liquid-liquid phase separation at high concentrations and low temperatures *(6–8)*. Moreover, the C-terminal domain of RNA Polymerase II (Pol II), itself an LCD, can be incorporated into FET LCD hydrogels in a phosphorylation-regulated manner *(9)*. FET LCDs were also reported to undergo phase separation in live cells upon overexpression *(7, 10)*.

However, there are stark differences between *in vivo* physiological conditions and those used for *in vitro* or overexpression studies. Indeed, temperature, protein concentration, purity and microenvironment, may all significantly affect the behavior of LCDs. There is also a vigorous debate over whether LCDs undergo cross-β polymerization or remain in a disordered conformation when interacting with partners *(6–8, 10–16)*. From the perspective of elucidating how TFs work *in vivo*, an equally pressing unresolved mechanistic question concerns the dynamics and time scales governing LCD-LCD interactions that would allow TFs to function in rapid cellular processes. Selectivity and specificity of cognate LCD-LCD interactions are other important yet poorly understood features required for proper TF function *in vivo*. Thus far, sequence-specific LCD-LCD recognition codes have not been directly demonstrated, let alone understood at a mechanistic level in the *in vivo* context. Here, we have combined a variety of high-resolution imaging strategies including fluorescence recovery after photobleaching (FRAP) *(17)*, lattice light sheet microscopy *(18)*, 3D DNA fluorescence in situ hybridization (FISH) *(19)*, and live-cell single-particle tracking (SPT) *(20, 21)*, to probe the dynamic behavior of TF LCDs at target genomic loci under physiological conditions.

## Results

We first established proof-of-concept experiments using a synthetic Lac operator (LacO) array (~50,000 LacO repeats) integrated into the genome of human U2OS cells *(22)* that express various EYFP-tagged TF-LCDs fused to LacI (Fig. 1A). To probe potential sequence specific LCD-LCD interactions, we examined two distinct classes of LCDs: QGYS-rich LCDs from the FET family (FUS/EWS/TAF15), and a QGTS-rich LCD from Sp1 that is low in tyrosine (Table S1).

**Fig. 1.**
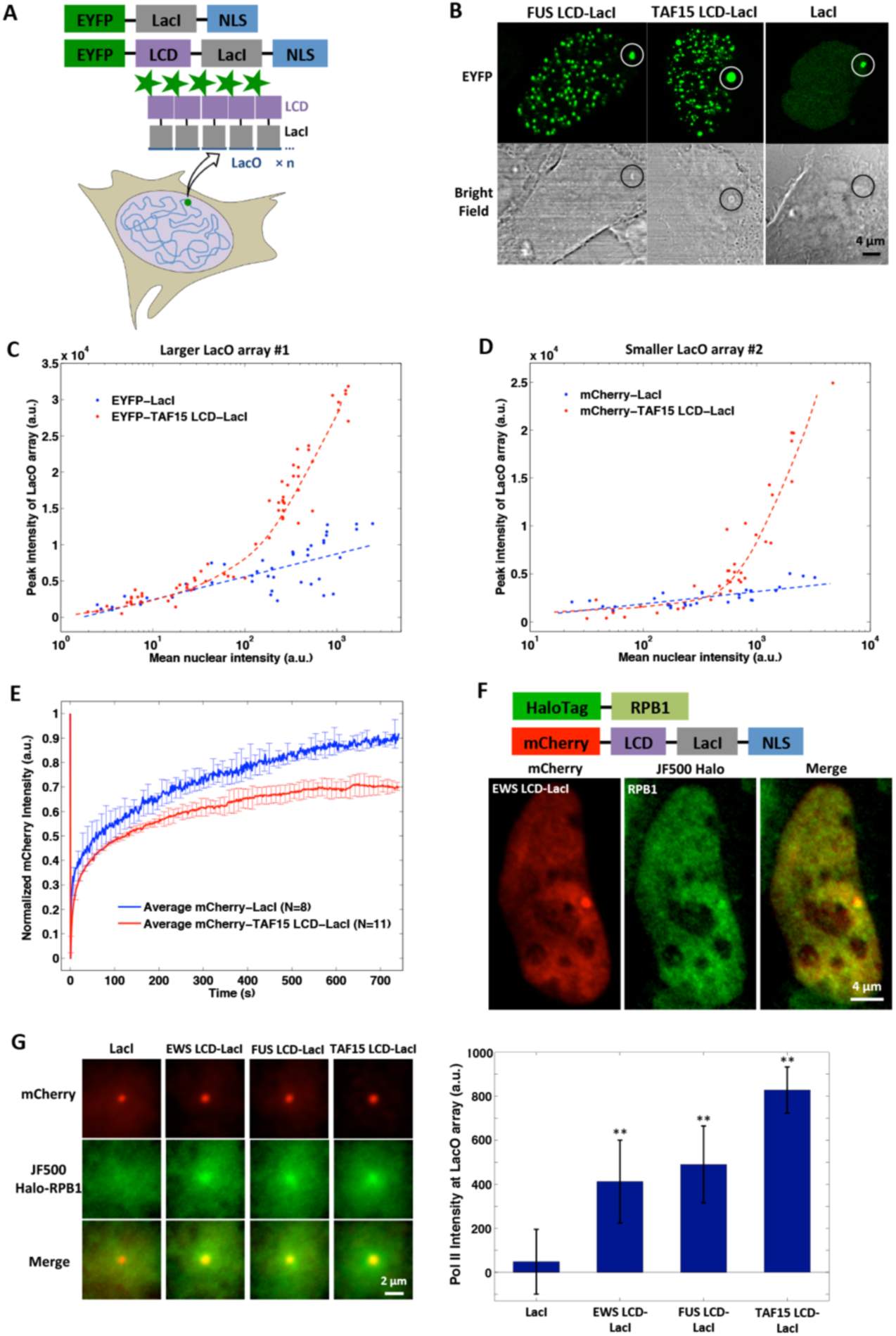
A LacO array can mediate the formation of an LCD hub in live cells, which involves extensive LCD self-interaction and recruits RNA Pol II. (**A**) Schematic for a LacO array (n ≈ 5 × 10^4^ for array #1, n ≈ 1.5 × 10^4^ for array #2) in the U2OS genome nucleating an LCD hub when EYFP-LCD-LacI is transiently expressed. Alternatively, EYFP-LacI is expressed as a control. NLS: nuclear localization signal. (**B**) Confocal fluorescence and bright field images of LacO-containing U2OS cells where LacO arrays #1 (highlighted by circles) are bound by EYFP-labeled LCD-LacI or LacI. LCD-LacI but not LacI bound LacO arrays are visible in bright field images. (**C-D**) Peak fluorescence intensity of LacO array #1 (**C**) or #2 (**D**) bound by EYFP-labeled (**C**) or mCherry-labeled (**D**) TAF15 LCD-LacI or LacI as a function of mean nuclear fluorescence intensity, which is proportional to the protein expression level. Each dot represents one cell. (**E**) Averaged FRAP curves at LacO array #1 bound by mCherry-labeled TAF15 LCD-LacI or LacI. Error bars represent standard deviations (SD). (**F**) (Upper) Schematic of the proteins expressed in the LacO-containing U2OS line. (Lower) Confocal fluorescence images show that Halo-RPB1 (labeled with 200 nM Halo ligand JF500, green) is enriched at LacO array #2 bound by mCherry-EWS LCD-LacI (red). (**G**) (Left) Averaged Halo-RPB1 images at LacO array #2 bound by mCherry-labeled LacI, EWS LCD-LacI, FUS LCD-LacI, or TAF15 LCD-LacI (N=55, 69, 81 or 143). (Right) Fluorescence intensity of Halo-RPB1 at the LacO array center in the average images after subtraction of nuclear Halo-RPB1 background (see supplemental methods). **: statistically significant increase compared to the LacI condition (p < 0.01, two-sample t-test). Error bars represent bootstrapped SD *(39)*.

As expected, the LacO array recruits a large number of EYFP-LCD-LacI molecules via targeted DNA binding, forming a concentrated local interaction hub in the nucleus. Intriguingly, LacO-associated hubs formed by LCD-LacI but not LacI are visible by bright field microscopy (Fig. 1B), suggesting that the refractive index and mass density of LCD-LacI hubs differ significantly from the surrounding nuclear environment.

Importantly, we found that both TAF15-LacI and Sp1-LacI give rise to much brighter and larger LacO-associated LCD “hubs” than LacI alone (Fig. 1C, S1A-B). The fluorescence intensity of LacO-associated LCD-LacI hubs increases with the TF nuclear concentration much faster than LacI hubs (Fig. 1C, S1B), suggesting increased binding valency of the LacO array for LCD-LacI that is contributed by extensive LCD self-interactions. Similarly, smaller LacO arrays containing significantly fewer (~15,000) LacO repeats also nucleate LCD self-interactions (Fig. 1D, S1C).

Moreover, LCD-LacI but not LacI alone can form hundreds of smaller puncta throughout the nucleus once its intra-nuclear concentration reaches a certain threshold (Fig. 1B, S1D-E). Interestingly, LCDs can form intra-nuclear puncta in some cases even without being fused to a DNA binding domain such as LacI (Fig. 3A, lower). These results suggest that LCD-LCD interactions can promote self-assembly of LCD hubs upon overexpression without the assistance from DNA *(7, 10)*.

In addition, FRAP dynamics of LCD-LacI at the LacO array was also significantly different from that of LacI (Fig. 1E). Since diffusion contributes negligibly to the FRAP dynamics (Fig. S2A), such differences can be attributed to changes in binding reaction rates. Specifically, when we fit the FRAP curves with a reaction-dominant two-binding-state (fast and slow dissociation) model *(23)*, we found that fusing TAF15 or FUS LCD to LacI leads to more than 60% reduction of both fast and slow dissociation rate constants of LacI (Fig. S2B-E). This result suggests that at increased local TF concentrations, TF-LCD hubs driven by LCD-LCD interactions stabilize TF binding to its cognate genomic site via multivalent contacts.

Having demonstrated homotypic LCD self-interactions, we next investigated the potential role of heterotypic LCD-LCD interactions in hub formation. First, we tested whether TF-LCD hubs can interact with RNA Pol II *in vivo* by using a LacO-containing U2OS line in which we replaced the endogenous RPB1 (major and catalytic subunit of Pol II) with an α-Amanitin-resistant Halo-tagged RPB1 *(24)*. We subsequently labeled the cells with a fluorescent HaloTag ligand and visualized Pol II distribution *in vivo*. We found that mCherry-FET-LacI expression mediates significant enrichment of Pol II in hubs compared to background levels recorded using LacI alone (Fig. 1F-G, S3). While recapitulating the *in vitro* incorporation of Pol II into LCD hydrogels *(9)*, these experiments go one step further and suggest that LCD hub formation can facilitate the recruitment of the general transcription machinery *in vivo* – a key step towards transactivation.

Next, we probed the sequence selectivity of various classes of LCD to interact with each other. To this end, we co-expressed both EYFP-LCD-LacI and mCherry-LCD in LacO-containing cells. mCherry-LCD lacking a DBD becomes enriched at the array only when it can interact with the co-expressed LCD that is fused to LacI (Fig. 2A, S4A). We note that the array can enrich mCherry-LCD over a wide range of expression levels. The EWS-LacI bound LacO array also enriches endogenous EWSR detected by immunofluorescence (Fig. S4B). Therefore, mCherry-LCD enrichment at the array is most likely due to specific LCD-LCD interactions rather than potential nonspecific overexpression artifacts. Using this two-color imaging assay, we confirmed homotypic self-interactions of all tested LCDs (from FET family and Sp1). Intriguingly, while all 3 FET LCDs interacted among themselves, none of them interacted with Sp1-LCD (Fig. 2A-B), suggesting that LCD interactions exhibit strong sequence specificity that is likely an essential feature underlying combinatorial TF regulation of gene expression.

**Fig. 2.**
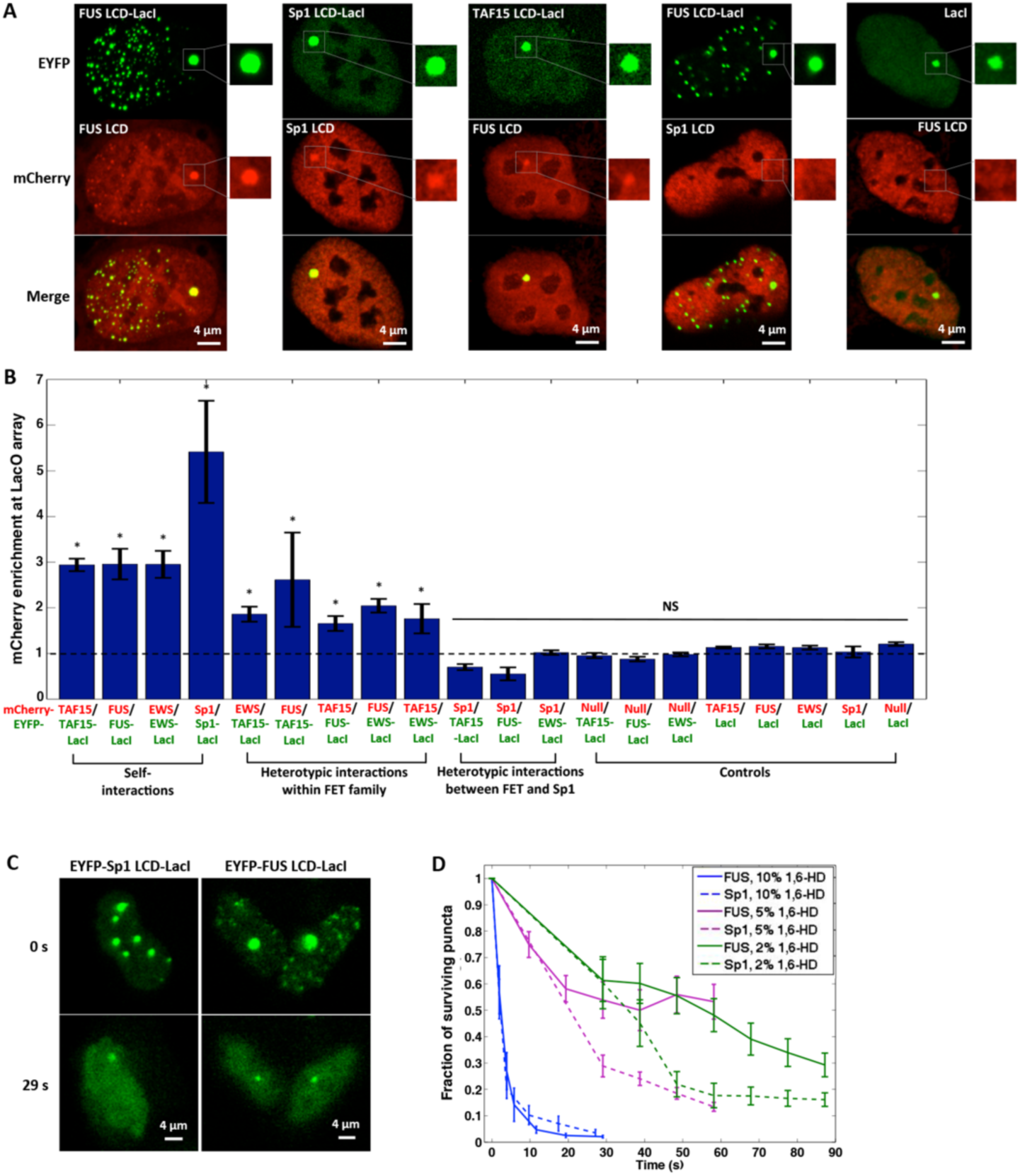
LCD hub formation involves selective protein-protein interactions, which can be disrupted by 1,6-HD with sequence-dependent sensitivity. (**A**) Confocal fluorescence images of U2OS cells containing LacO array #1 that co-express various combinations of mCherry-LCD and EYFP-LCD-LacI. The region surrounding the LacO array is zoomed in. (**B**) Quantification of the enrichment of mCherry-LCD (font color: red) at the LacO array #1 bound by various EYFP-labeled LCD-LacI fusion proteins (font color: green), calculated as the peak mCherry fluorescence intensity at the array divided by the average intensity immediately surrounding the array (see Fig. S4A). Null: mCherry not fused to any LCD. An mCherry enrichment at the array above 1 suggests LCD-LCD interactions. *: statistically significant difference above 1 (p < 0.05, one-sample t-test). NS: not significant difference above 1. Error bars represent standard errors (SE). (**C**) Fluorescence images of FUS and Sp1 LCD hubs before (0 s) and after (29 s) addition of 10% 1,6-HD. (**D**) Number of nuclear puncta formed by FUS or Sp1 LCD surviving over time upon addition of 1,6-HD at different concentrations. Error bars represent SE.

To better understand the nature of LCD-LCD interactions, we treated cells with 1,6-hexanediol (1,6-HD), an aliphatic alcohol known to dissolve various intracellular membrane-less compartments and FUS hydrogels *in vitro* through disruption of hydrophobic interactions *(25–27)*. We observed that both FUS and Sp1 LCD hubs rapidly disassemble within 30 sec when exposed to 10% 1,6-HD. The LacO associated LCD induced hub shrank to a size comparable to the array bound by LacI alone, while all the nuclear puncta not associated with LacO disappeared (Fig. 2C). We also found that 2,5-hexanediol (2,5-HD), a less hydrophobic derivative of 1,6-HD that barely melts FUS hydrogels *in vitro (25)*, disrupts LCD hubs less efficiently in live cells (Fig. S4C-D). This strong correlation between hydrophobicity of hexanediols and LCD hub melting suggests that these aliphatic alcohols directly influence LCD-LCD interactions by disrupting key hydrophobic contacts, rather than by affecting nonspecific global cell physiology. Our *in vivo* results also closely mirror *in vitro* hydrogel studies using these disrupting agents *(25)*.

Intriguingly, Sp1 LCD hubs were disrupted significantly faster and more extensively than FUS LCD hubs with 2% or 5% 1,6-HD (Fig. 2D). Thus, while a combination of intermolecular forces may contribute to LCD hub formation, our results indicate that hydrophobic interactions might be more sensitive to disruption and play a more dominant role in Sp1 LCD self-interactions than FUS LCD, consistent with the Sp1 LCD containing hydrophobic residues sparsely interspersed amongst Q repeats *(28)*. The differential sequence dependence of LCD-LCD interactions revealed by 1,6-HD treatment may be correlated to the selectivity of homo-and heterotypic LCD interactions observed above.

To study the dynamics of protein-protein interactions between LCD pairs, we co-expressed EYFP-LCD-LacI and Halo-LCD in the LacO-containing U2OS cells, and performed single particle tracking (SPT) of Halo-LCD to measure residence times (RTs) of LCD-LCD interactions within the LacO-associated hub (Fig. 3A, upper). For all LCDs tested, RTs resulting from self-interactions fell in the range of 11~33 sec (Fig. 3B). Interestingly, when EYFP-LCD and Halo-LCD from the FET family were co-expressed at high levels, they spontaneously formed hubs unaffiliated with the array that resemble intranuclear puncta (Fig. 3A, lower). These non-array hubs bind Halo-LCD via homo-or heterotypic interactions with even shorter RTs (7~10 sec). As expected, the Sp1 LCD that failed to interact with the FUS LCD, had an RT at the non-array FUS LCD hubs of <1 sec (Fig. 3B). The fact that RTs of many LCDs in self-aggregated hubs unaffiliated with genomic DNA are substantially shorter than in hubs formed at the LacO array suggests that TF-DNA interactions that maintain a high local concentration of TF-LCDs contribute to stabilizing LCD-LCD interactions and vice versa. Together, these findings reveal the rapid, reversible and interdependent nature of LCD-LCD and TF-DNA interactions as well as their propensity to form local high concentration hubs that likely stabilize multi-component complexes, *e.g*. transcription preinitiation complex, a prerequisite for transactivation.

**Fig. 3.**
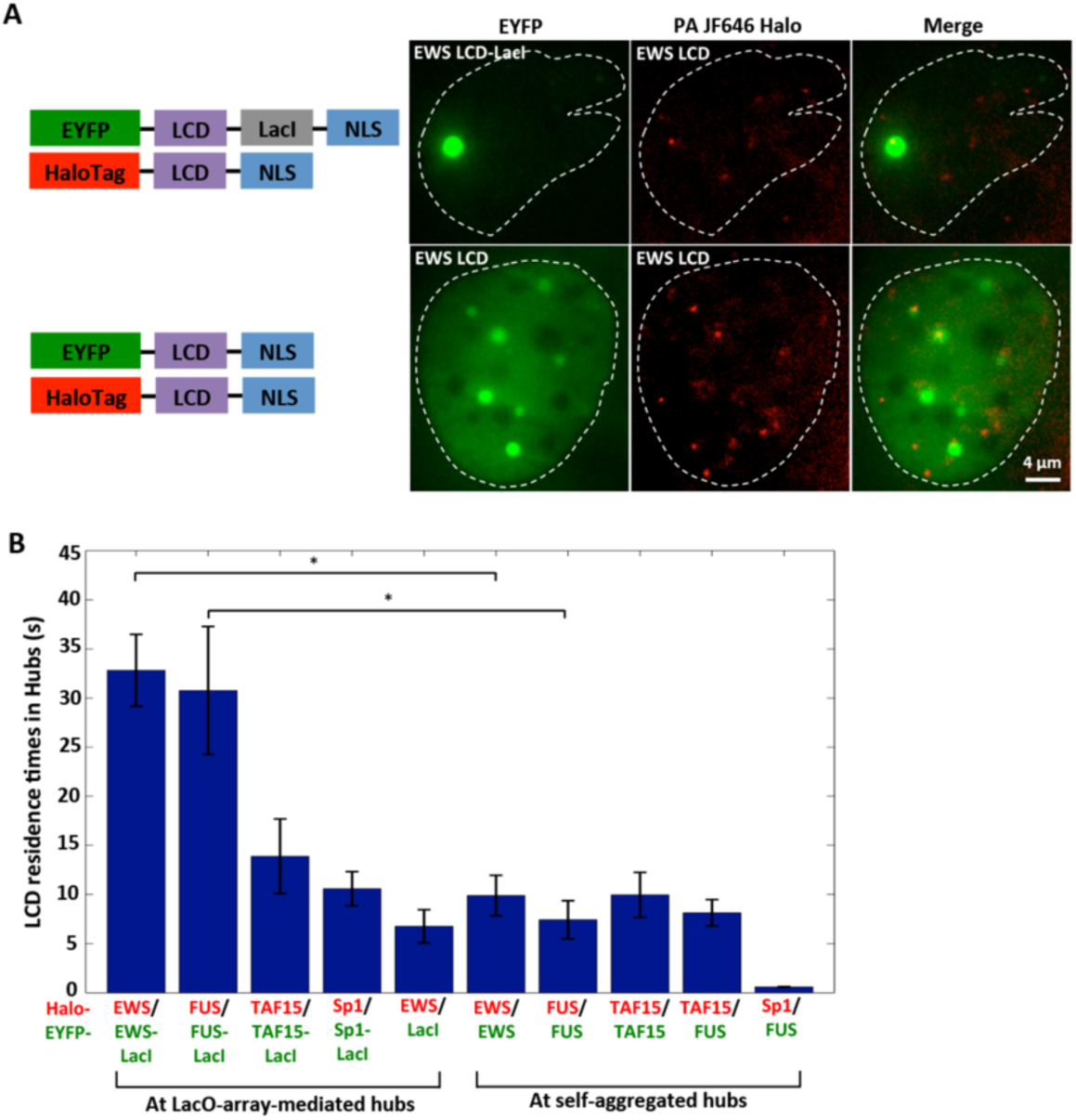
LCD-LCD interactions involved in hub formation are highly dynamic. (**A**) Snapshots of a two-color SPT movie simultaneously imaging EYFP-labeled (green) EWS LCD-LacI (upper, forming a LacO-associated LCD hub) or EWS LCD (lower, forming self-aggregated LCD hubs not affiliated with the LacO array) and Halo-tagged EWS LCD (2 nM PA-JF646 labeled, red) in U2OS cells containing LacO array #1. A white dashed contour outlines the cell nucleus. We imaged the hubs in the EYFP channel (green), and tracked individual Halo-EWS LCD molecules with an acquisition time of 500 ms in the PA-JF646 channel (red). (**B**) Residence times of LCD (font color: red) bound at the LacO-array-associated LCD hub or self-aggregated LCD hubs not affiliated with the array (font color: green). *: p < 0.05, two-sample t-test. Error bars represent SE.

Having unmasked the sequence specificity and dynamic nature of LCD-LCD interactions using synthetic LacO arrays in living cells, we next tested LCD behavior at native GGAA microsatellites (>20 GGAA repeats) in the Ewing’s sarcoma cell line A673 *(29–31)*. These cancer-derived cells have suffered a chromosomal translocation t(11;22)(q24;q12) producing a fusion oncogene, *EWS/FLI1*, that encodes a potent TF consisting of a trans-activating LCD from EWS and the DBD from FLI1 that targets GGAA sequences (Fig. 4A).

**Fig. 4.**
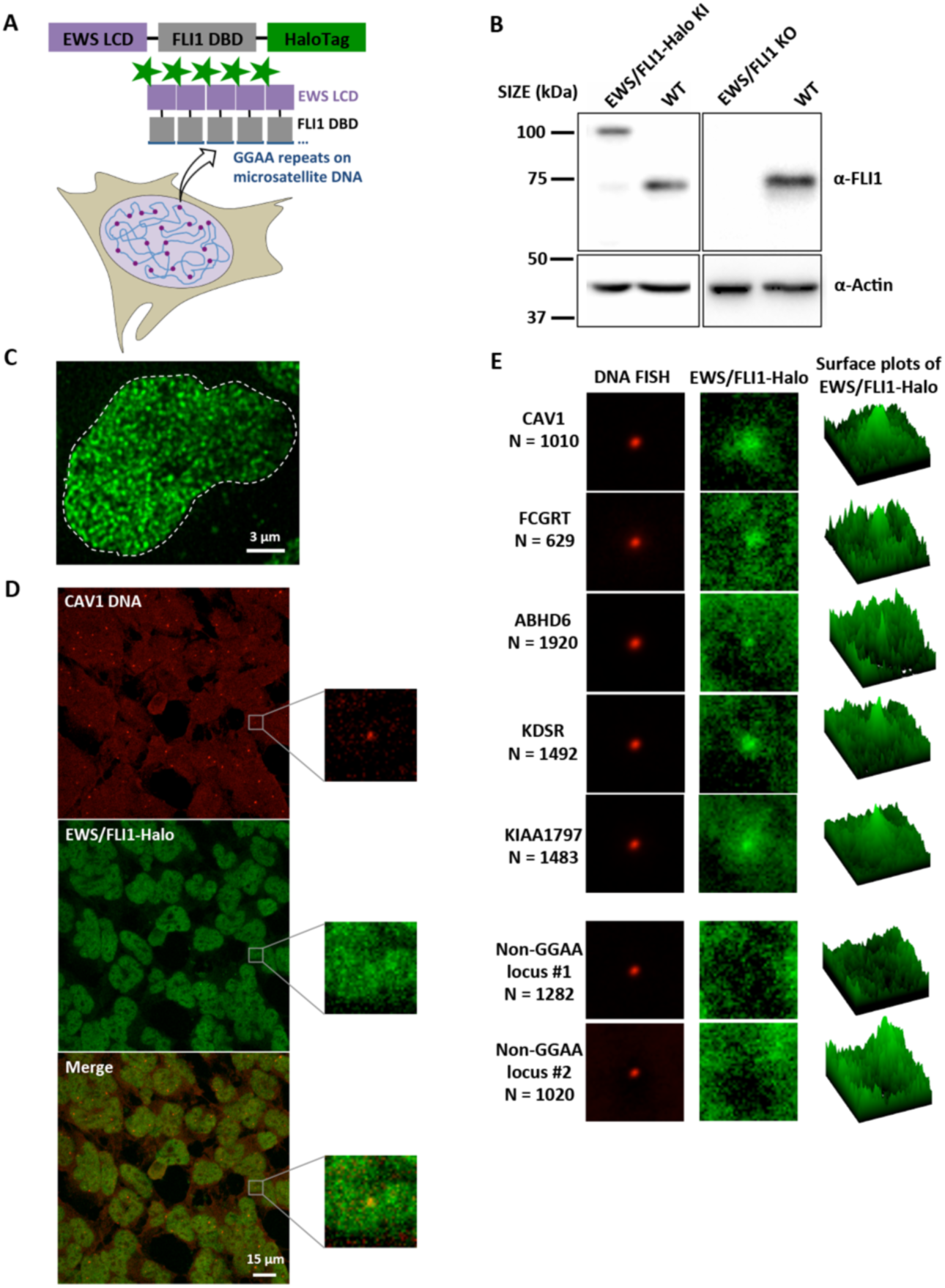
Combined DNA FISH and EWS/FLI1-Halo imaging show that endogenous EWS/FLI1 forms hubs at GGAA-microsatellites. (**A**) Schematic for GGAA-microsatellites in the A673 genome nucleating hubs of endogenously Halo-tagged EWS/FLI1. (**B**) Western blot of EWS/FLI1 and Actin (normalization control) from clonal EWS/FLI1-Halo knock-in (KI), wild-type, and clonal EWS/FLI1 knockout (KO) A673 lines. (**C**) z-projected 3D image of endogenous EWS/FLI1-Halo in an A673 cell nucleus (stained with 200 nM Halo ligand JF549) taken on the lattice light sheet microscope. (**D**) Confocal fluorescence images of 3D DNA FISH targeting GGAA-microsatellite-adjacent *CAV1* gene (enhanced Cy5 labeled, red) and endogenous EWS/FLI1-Halo (JF549 labeled, green). The region surrounding one particular *CAV1* locus is zoomed in. EWS/FLI1-Halo enrichment at the locus is visible but buried in high background. (**E**) Averaged two-color images of five GGAA-microsatellite-adjacent gene loci (*CAV1, FCGRT, ABHD6, KDSR, KIAA1797*) and two gene loci not containing a GGAA microsatellite (Non-GGAA #1 targeting *ADGRA3* gene and #2 targeting *REEP5* gene). The right column shows average surface plots of EWS/FLI1-Halo.

In order to visualize the behavior of endogenously expressed EWS/FLI1, we fused a HaloTag to its DBD using CRISPR/Cas9-mediated genome editing of A673 cells (Fig. 4A-B, S5A) *(32)*. This knock-in strategy allowed us to image fluorescently tagged endogenous EWS/FLI1 at its normal expression levels (Fig. 4B). This is essential as LCDs tend to self-aggregate and behave aberrantly upon overexpression. To ensure that Halo-tagging does not disrupt transactivation functions of EWS/FLI1, we confirmed that EWS/FLI1-Halo activates a luciferase reporter construct containing a GGAA-microsatellite-driven promoter *(30)* as efficiently as WT EWS/FLI1 (Fig. S5C). More importantly, using the gold standard neoplastic transformation assay *(33)*, we confirmed that the EWS/FLI1-Halo knock-in A673 cells form colonies in soft agar much like the WT A673, albeit, less efficiently (Fig. S5D-E).

We next performed high-resolution lattice light sheet microscopy and found that EWS/FLI1 forms many small interaction hubs (>1000 per nucleus) in the nucleus (Fig. 4C). The detected number of intra-nuclear hubs has the same order of magnitude as the number of EWS/FLI1-bound GGAA microsatellites across the human genome (~6000) estimated by ChIP-seq and bioinformatics analyses *(34)*. To examine the spatial relationship between EWS/FLI1 hubs and GGAA microsatellites, we performed simultaneous confocal imaging of EWS/FLI1-Halo and 3D DNA FISH targeting genes adjacent to GGAA microsatellites that are regulated by EWS/FLI1 (Fig. 4D), including *CAV1, FCGRT, ABHD6, KDSR* and *KIAA1797 (30, 35)*. Although EWS/FLI1 enrichment is detected at many single loci of these genes, the crowded distribution of intra-nuclear EWS/FLI1 hubs makes it difficult to clearly visualize EWS/FLI1 enrichment at single target loci. By recording images of ~1000 loci for each gene, the signal to noise ratio is significantly improved to reveal specific EWS/FLI1 enrichment at GGAA repeats, while no enrichment was seen at non-GGAA gene loci (Fig. 4E). These results indicate that EWS/FLI1 forms hubs at endogenous GGAA microsatellite DNA elements.

We previously observed that the formation of an LCD interaction hub slows down dissociation of LCD-LacI from the LacO array (Fig. 1E, S2E). If LCD-LCD interactions are also involved in EWS/FLI1 hub formation at GGAA microsatellites, we expect the residence time of EWS/FLI1 within GGAA-affiliated hubs to be longer than outside hubs. We stained the EWS/FLI1-Halo knock-in A673 cells with two fluorescent ligands *(36, 37)*: high-concentration JF549 staining allows visualization of EWS/FLI1 hubs in the cell nucleus, while low-concentration PA-JF646 staining allows real-time tracking of individual EWS/FLI1 molecules (Fig. 5A). SPT revealed the average RTs of EWS/FLI1 in and outside the GGAA-affiliated hubs to be 90 sec and 16 sec, respectively (Fig. 5B, S6A-B). The fact that EWS/FLI1 binds to GGAA repeats for significantly longer time suggests that EWS LCD-LCD interactions are likely involved in the formation of the GGAA-affiliated hubs. Therefore, very likely both LCD-LCD interactions and DNA binding to GGAA-repeats work together to stabilize hub formation much as we observed for LCD-LacI at the LacO array.

**Fig. 5.**
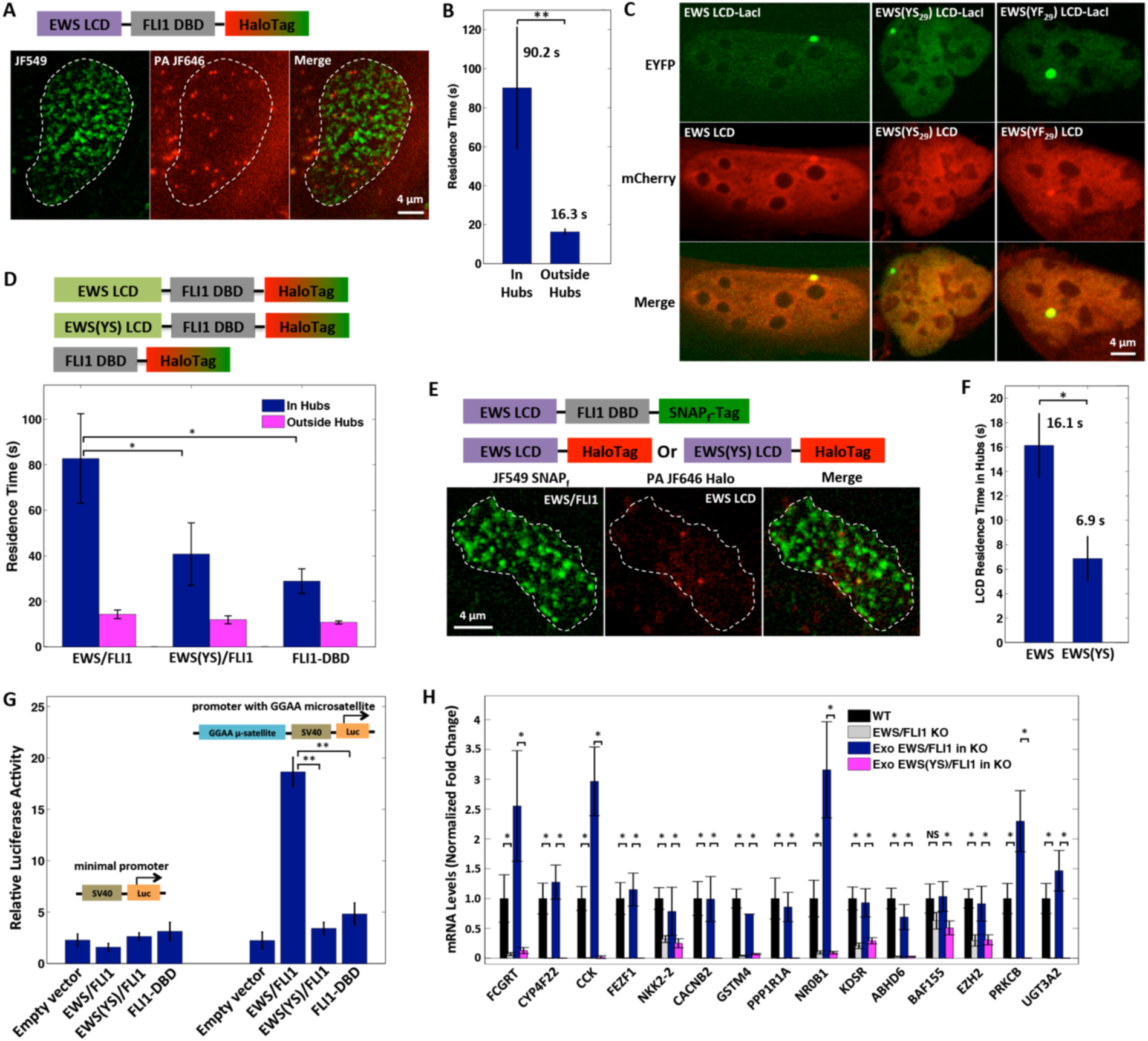
Dynamic LCD-LCD interactions occur at GGAA microsatellites, which stabilize EWS/FLI1 binding and drive its transactivation function. (**A**) Snapshots of an SPT movie imaging endogenous EWS/FLI1-Halo labeled with two Halo ligands, JF549 (200 nM) and PA-JF646 (20 nM). We imaged the EWS/FLI1-Halo hubs in the JF549 channel (green), and tracked individual EWS/FLI1-Halo molecules in and outside the hubs in the PA-JF646 channel (red). (**B**) Residence times of EWS/FLI1 bound in hubs are longer than outside hubs as determined by SPT (p < 0.01, two-sample t-test). Error bars represent SE. (**C**) EWS LCD is enriched at the LacO array #1 bound by EWS LCD-LacI, but EWS(YS_29_) LCD is not recruited to the array by EWS(YS_29_) LCD-LacI. However, EWS(YF_29_) LCD is recruited to the array by EWS(YF_29_) LCD-LacI. (**D**) (Upper) Schematic of proteins transiently expressed in EWS/FLI1 KO A673 cells: Halo-tagged EWS/FLI1, EWS(YS)/FLI1 or FLI1 DBD. (Lower) Residence times of EWS/FLI1 and its variants binding in and outside their hubs as determined by SPT. *: p < 0.05, two-sample t-test. Error bars represent SE. (**E**) Snapshots of an SPT movie simultaneously imaging SNAPf-tagged EWS/FLI1 (200 nM JF549 labeled, green) and Halo-tagged EWS or EWS(YS) LCD (20 nM PA-JF646 labeled, red) in EWS/FLI1 KO A673 cells. Individual LCD-Halo molecules were tracked with the strategy described in (A). (**F**) Residence times of EWS bound at EWS/FLI1 hubs are longer than EWS(YS) LCD as determined by SPT (p < 0.05, two-sample t-test). Error bars represent SE. (**G**) Luciferase assay shows that EWS/FLI1 but not EWS(YS)/FLI1 or FLI1 DBD transactivates a GGAA-microsatellite-driven reporter (p < 0.01, two-sample t-test). Error bars represent SE. (**H**) RT-qPCR shows downregulation of GGAA-microsatellite-associated EWS/FLI1 target genes in A673 cells upon EWS/FLI1 KO. Stable expression of exogeneous (Exo) EWS/FLI1, but not of the mutant EWS(YS)/FLI1, rescues the expression defect in EWS/FLI1 KO A673 cells. For each target gene, the mRNA level was normalized using 5 different invariant genes (Fig. S9A) and graphed as a fold change relative to the mRNA level present in the WT A673 line (set to 1). *: p < 0.05, two-sample t-test. NS: not statistically significant. Error bars represent SD.

To confirm that hub formation in this native setting is dependent on EWS-LCD, we determined how mutations in the LCD might affect RTs of EWS/FLI1. We started by replacing different numbers (m = 3, 7, 10, 17 or 29) of tyrosines (Y) in the EWS LCD (residues 47–266 of EWS) with serines (S), and testing the self-interaction capability of mutant LCDs (EWS(YS_m_)) using the LacO array assay established earlier. As previously shown, when we co-expressed EYFP-EWS-LacI and mCherry-EWS, the mCherry signal became enriched at the LacO array due to EWS LCD self-interaction. Interestingly, when we replaced WT EWS in both fusion proteins with EWS(YS_m_), mCherry enrichment at the array progressively decreased with increasing number of Y-to-S mutations (Fig. S7A), and vanished for EWS(YS_29_) where all the tyrosines are replaced (Fig. 5C). Similarly, we found that EWS(YS_29_) does not interact with WT EWS (Fig. S7B-C). By contrast, a mutant replacing all 29 tyrosines with phenylalanine (EWS(YF_29_)) retains hub formation activity with itself and with WT EWS (Fig. 5C, S7D), suggesting that aromatic amino acids and hydrophobic contacts represent a major driver of EWS LCD-LCD interactions.

Next, we probed the effects of mutations that disrupt LCD hub formation on RTs of EWS/FLI1. In order to examine behaviors of EWS/FLI1 variants in A673 cells without interference of endogenous EWS/FLI1, we generated an EWS/FLI1 knockout A673 line using CRISPR/Cas9-mediated genome editing (Fig. 4B, S5B) and verified that transiently and moderately re-expressed EWS/FLI1-Halo in the knockout line exhibited binding dynamics comparable to that of endogenous EWS/FLI1-Halo (Fig. 5B&D, S6C). We then transiently expressed similar levels of a Halo-tagged LCD deletion mutant (FLI1 DBD) or a 37-residue Y-to-S mutant (EWS(YS)/FLI1). Both mutants still displayed some hubs in the nucleus, but they are significantly diminished and SPT revealed that their in-hub RTs become significantly reduced (by 51–65%) relative to WT EWS/FLI1 while their outside-hub RTs remain largely unchanged (Fig. 5D). Together, these results confirm that LCD-LCD interactions drive the formation of EWS hubs at GGAA microsatellites.

We previously showed that the fluorescence intensity of LacO-associated LCD-LacI hubs increases with the TF nuclear concentration much faster than LacI hubs due to extensive LCD-LCD interactions (Fig. 1C-D, S1B-C). Similarly, when transiently expressing EWS/FLI1-Halo or FLI1 DBD-Halo in U2OS cells, we found the fluorescence intensity of EWS/FLI1 hubs increases faster than FLI1 DBD alone as a function of TF concentration (Fig. S8). This finding further confirms that LCD-LCD interactions are involved in GGAA-affiliated EWS/FLI1 hubs.

To measure the dynamics of just the protein-protein interactions occurring within the EWS LCD hubs, we transiently expressed SNAPf-tagged EWS/FLI1 and Halo-tagged EWS in the EWS/FLI1 knockout line, and labeled both fusion proteins using fluorescent ligands with distinct emission spectra *(36, 37)*. While EWS/FLI1-SNAPf forms hubs at GGAA microsatellites via protein-DNA binding, EWS LCD-Halo, which does not interact with DNA, binds to EWS/FLI1 hubs only via protein-protein interactions. We visualized EWS/FLI1 hubs and simultaneously tracked individual EWS LCD molecules that bind to the hubs (Fig. 5E). SPT revealed the average RT of EWS LCD in EWS/FLI1 hubs to be 16 sec, suggesting LCD-LCD interactions are highly dynamic (Fig. 5F). As expected, the mutant EWS(YS) LCD has a significantly shorter RT (~7 sec) at EWS/FLI1 hubs, consistent with its diminished interaction with EWS LCD.

Finally, we tested whether LCD-LCD interactions influence EWS/FLI1 functions. Importantly, we found that whereas EWS/FLI1 efficiently induces gene activation at a GGAA microsatellite in a luciferase assay, the mutant EWS(YS)/FLI1 and the FLI1 DBD do not (Fig. 5G). We further engineered the EWS/FLI1 knockout A673 line to stably express EWS/FLI1 or EWS(YS)/FLI1, performed RT-qPCR to measure the expression levels of GGAA-microsatellite-associated EWS/FLI1 target genes. As expected, we found that expression of EWS/FLI1, but not EWS(YS)/FLI1, specifically rescues the gene expression defect in the knockout line, indicating that EWS LCD-LCD interactions are required for transactivation (Fig. 5H). Moreover, the knockout line stably expressing EWS(YS)/FLI1 does not form colonies in soft agar like wild-type A673 (Fig. S9B). This demonstrates that EWS LCD-LCD interactions are required for oncogenic transformation. Taken together with previously published RT-qPCR and RNA-seq data in mesenchymal stem cells showing the important role of EWS LCD in inducing expression of GGAA-microsatellite-associated genes *(38)*, our results suggest that the formation of EWS LCD-dependent hubs is essential for EWS/FLI1 to activate transcription and drive oncogenic gene expression programs in Ewing’s sarcoma.

## Discussion

Through imaging TF-LCD interactions in live cells, our findings offer a powerful complement to pioneering *in vitro* studies that provided the first clues about LCD interactions *(9)*. Importantly, to the extent that one can make comparisons between hydrogels and intracellular LCD hub formation, many aspects of FET-LCD function uncovered *in vitro* are born out when probed under physiological settings in live cells. In addition, single-molecule live-cell imaging revealed several new aspects of LCD-driven interactions. Most striking is the highly dynamic and sequence-specific nature of LCD interactions as they form local high concentration hubs that drive transactivation. We were also especially intrigued by the formation of LCD-dependent hubs throughout the nucleoplasm that are not associated with cognate genomic DNA. These LCD-LCD interaction driven puncta, especially their sensitivity to 1,6-HD and highly dynamic nature (RTs of 7–10 sec), provide new insights into mechanisms governing transactivation.

Although our studies were not designed to address the structure and nature of LCD-driven phase separated compartments, under overexpression conditions we detected what appears to be phase separation (*i.e*., local changes in refractive index). Although we did not obtain direct evidence for phase separation for TF (*i.e*. EWS/FLI1) hubs formed under endogenous expression levels, we have detected functionally relevant LCD-LCD interactions involved in these TF hubs. Given the short RTs and highly transient nature of LCD-LCD transactions we measured, LCD-dependent transactivation can apparently occur in hubs formed within a broad range of local TF concentrations and time scales – from hyper rapid non-specific LCD-LCD and TF-DNA binding events (0.1–1 sec) all the way to relatively stable aggregates (mins). Both the composition and diversity of LCDs in hubs and their interaction specificity could influence the range of their operational concentrations and their potential for phase separation and/or polymer formation. New insights regarding the rapid binding dynamics and functional importance of TF-LCDs (*i.e*. LCD in the oncogenic EWS/FLI1) suggest that understanding these mechanisms may also enhance our ability to develop novel strategies to modulate gene expression in certain disease settings. Finally, although we examined a small subset of TF-LCDs, the fundamental principles that we have uncovered about the dynamics and mechanisms driving LCD-LCD transactions may be applicable to other classes of regulatory proteins and biomolecular interactions occurring in a variety of cell types.

## Acknowledgments

We thank Stephen Lessnick and Masato Kato for helpful discussions and generously providing reagents, Qijian Gan and Anders Hansen for providing codes to analyze imaging data, James Bosco and Pranav Sharma for help with molecular cloning and James Goodrich and members of Tjian and Darzacq labs for critical reading of the manuscript. This work was supported by California Institute of Regenerative Medicine grant LA1-08013 (to X.D.), the National Institutes of Health grants UO1-EB021236 and U54-DK107980 (to X.D.), and the Howard Hughes Medical Institute (to Z.L., L.L. and R.T.).

